# When paleontology meets genomics: complete mitochondrial genomes of two saber-toothed cats’ species (Felidae: Machairodontinae)

**DOI:** 10.1101/2023.09.14.557696

**Authors:** Igor Henrique Rodrigues-Oliveira, Iuri Batista da Silva, Renan Rodrigues Rocha, Rafael Augusto Silva Soares, Fabiano Bezerra Menegídio, Caroline Garcia, Rubens Pasa, Karine Frehner Kavalco

## Abstract

Within the Machairodontinae subfamily, commonly referred as saber-toothed cats, it is worth noting that only two species, namely *Homotherium latidens*, recognized as the scimitar-toothed cat, and *Smilodon populator*, renowned as the saber-toothed tiger, possess partial mitochondrial genomes accessible in the NCBI database. These sequences, however, do not include the mitogenome control region (D-loop) and have several gaps in their genes, including protein-coding genes (PCGs) that are widely used in phylogenetic analysis. In this study, we aimed to obtain a complete assembly of the mitogenomes of these two species from next-generation sequencing data available at NCBI’s SRA. The *de novo* assemblies showed complete mitogenomes with 17,323bp (*H. latidens*) and 16,769 bp (*S. populator*), both with 13 PCGs, 22tRNAs, two rRNAs and the D-loop (Control region), with all genes following the standard order and position of most vertebrate mitogenomes. Our phylogeny and molecular dating, despite being generally very similar to previous studies, reveals an earliest divergence between North American and North Sea *H. latidens* specimens which may be related to an Early Pleistocene migration across Beringia.

## 1 Introduction

The saber-toothed cats of the Machairodontinae subfamily were among the apex predators of the Miocene to Pleistocene epochs, with an Early Miocene (around 18ma) divergence between Smilodontini and Homotherini tribes ^1^. Only a few species of the group survived until the Late Pleistocene, in general belonging to *Smilodon* and *Homotherium* genera. For *Smilodon*, two different species in the Late Pleistocene are recognized: *Smilodon populator* from east of the Andean South America and *Smilodon fatalis* from west of the Andean South America, Central America, and North America ^2^, with an isolate record from Uruguay in the east of the Andean South America ^3^. For *Homotherium*, however, the number of Late Pleistocene species has been the subject of debate in recent years. Traditionally Late Pleistocene specimens of *Homotherium* from Eurasia are assigned to *Homtherium latidens* while in North America to *Homotherium serum* ^4^. However, morphological and mitochondrial DNA data were used to propose that all Late Plesitocene Holarctic specimens consisted of a single species: *H. latidens* ^1,5,6^. In one of these studies, Paijmans et al. ^1^ described the partial mitochondrial genome of one *S. populator* and three *H. latidens* (two from North America) and used the assembled sequences to make phylogenetic inferences with other Felidae mitogenomes, after aligning these sequences and remove missing data, resulting in a 6,649 bp alignment with five partitions calculated by PartitionFinder. Besides presenting most of the mitogenome of these species, the assemblies of Paijmans et al. ^1^ lack the control region (D-loop) of the mitogenome and have several gaps in their genes, including protein-coding genes (PCGs). To improve the knowledge of the organization, composition and phylogeny of the mitochondrial genome of this group, we present the first assembly of the complete mitogenome of one specimen of *S. populator* and one *H. latidens* by a *de novo* method, as well as other two assemblies of *H. latidens* by a mapping method.

## 2 Results and Discussion

### 2.1 The complete mitochondrial genome of *S. populator* and *H. latidens*

We assembled the complete mitochondrial genome of *S. populator* and *H. latidens* by a *de novo* method using libraries available in the Sequence Read Archive (SRA) from the NCBI database. For *S. populator* we used the DNA-Seq library SRR13403298 from Westbury et al. ^7^ study and for *H. latidens* we used the SRR12354130 from Barnett et al. ^8^. The mitochondrial genome of *S. populator* ZMA20.042 presents 16,769 bp (Figure1a), in contrast with the Paijmans ^1^assembly which has 15,459 bp. The mitochondrial genome of *H. latidens* YG 439.38, on the other hand, presents 17,323 bp (Figure1b) in contrast to 15,471 bp from the same fossil in addition to 15,462 and 15,437 bp from other fossils assembled by Paijmans ^1^ . The maximum alignment score for the *S. populator* mitochondrial genome in the NCBI nt database was the complete mitochondrial genome of *Prionailurus bengalensis* (99% of query cover and 86.99% of identity), while for *H. latidens* was the *Prionailurus iriomotensis* mitochondrial genome (99% of query cover and 85.57% of identity). Since it was not possible to obtain a *de novo* assembly with the libraries of *H. latidens* specimens SP1007 and SP1714, we assembled the mitogenomes of these specimens by mapping the libraries against our *H. latidens* YG 439.38 assembly. The *H. latidens* SP1007 mitogenome presents 17,369 bp, with 33,17% of missing data (Figure1c) and the *H. latidens* SP1714 mitogenome presents 17,334 bp, with 4,77% of missing data (Figure1d). The GC content in *S. populator* ZMA20.042 mitogenome was 38.99% and in *H. latidens* YG 439.38 mitogenome was 37.74%. For *H. latidens* SP1007 and SP1714 the GC content was smaller due to the number of ambiguous nucleotides in the assembly, being 26.47% and 35.83% respectively. The main difference between ours and the Paijmans ^1^assemblies is the presence of the presence of the control region, also called D-loop, in our assemblies. Besides this, we also solved the gaps in the previous assemblies. For the *S. populator* ZMA20.042 assembly from Paijmans ^1^ few gaps could be observed in genes *COX2, ATP6, COX3, ND3, ND4, ND5* and *CYTB* (Figure1a). For *H. latidens* fossil YG 439.38 assembled by Paijmans ^1^ large gaps could be observed in genes *ND2, COX2, ATP8, ATP6, COX3, ND3, ND4, ND5* and *CYTB*, besides the complete absence of *ND4L* (Figure1b). Both species presented the standard gene order and composition of most vertebrate mitogenomes, with 22 tRNAs, 13 PCGs and two rRNAs ^9^. Among the PCGs only the *ND6* was found in the light strand, while for non-coding genes eight tRNAs were found in light strand (*tRNA-Gln, tRNA-Ala, tRNA-Asn, tRNA-Cys, tRNA-Tyr, tRNA-Ser, tRNA-Glu* and *tRNA-Pro*).

**Figure1.**
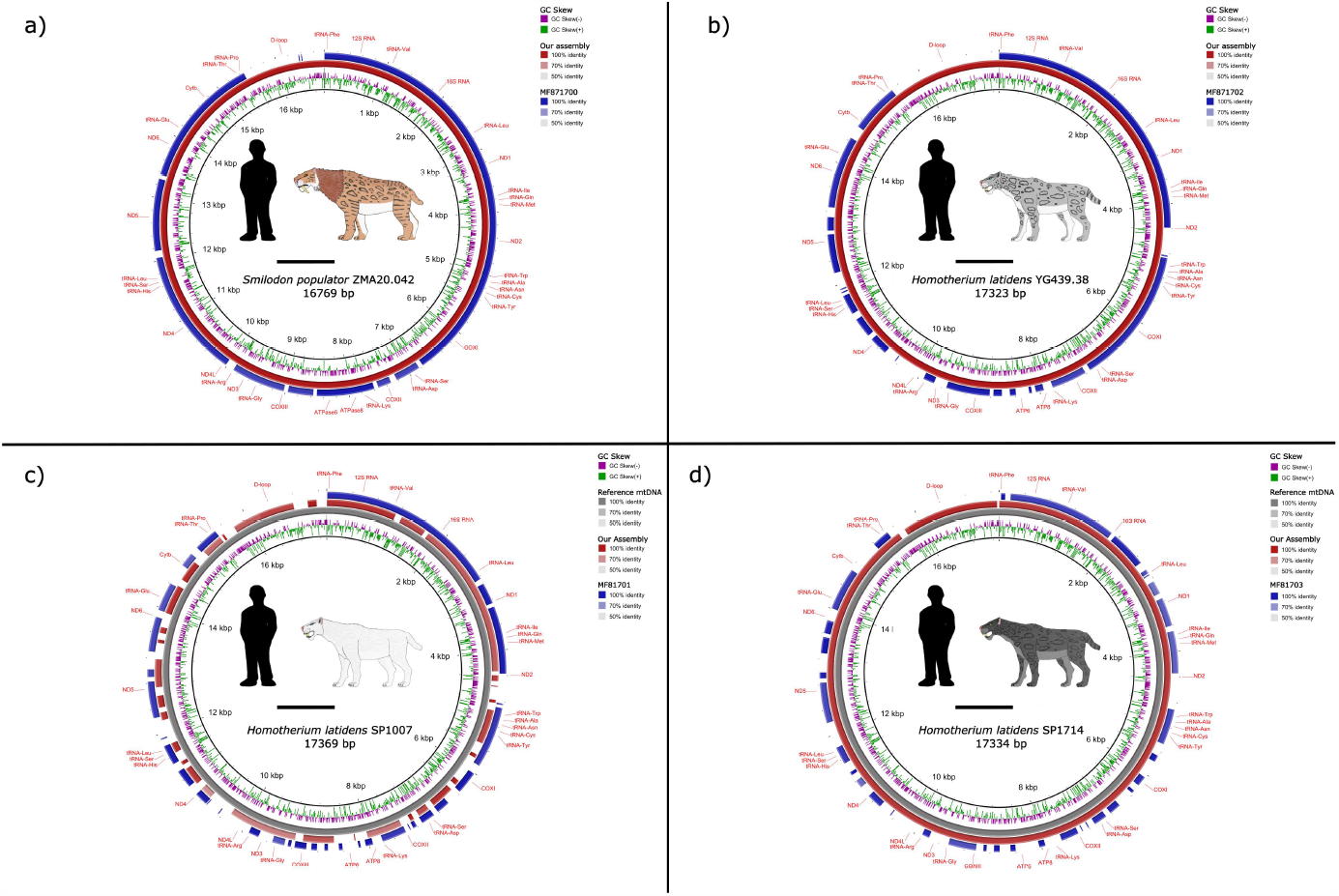
BLAST comparisons of our assemblies against assemblies of the same specimens in the NCBI nt database. (A) *Smilodon populator* ZMA20.042. (B) *Homotherium latidens* YG 438.38. (C) *Homotherium latidens* SP1007. (D) *Homotherium latidens* SP1714.

Considering only mitogenomes assembled by the *de novo* method (*S. populator* ZMA20.042 and *H. latidens* YG 439.38), base composition was 33.04% A, 27.97% T, 25.46% C and 13.53% G for *S. populator*, and 33.18%% A, 29.09% T, 24.95% C and 12.79% G for *H. latidens*. The 13 PCGs comprehends 11,339 bp of *S. populator* mitogenome, while the *12S rRNA* and *16S rRNA* occupy together 2,544 bp, the 22 tRNAs occupy 1,523 bp and the D-loop occupy 1,310 bp. For *H. latidens*, on the other hand, the 13 PCGs occupy 11,336 bp, the rRNAs (12S and 16S) occupy 2,543 bp, the 22 tRNAs occupy 1,516 bp and the D-loop occupy 1,856 bp of the complete mitogenome. It is remarkable that the mitogenomes of both saber-toothed cats have nucleotide composition and gene size very similar to each other and to the mitogenome of other felids, with most of the variation occurring in the mitogenome control region, as has been previously reported among felids mitogenomes ^10,11^. The control region of Felidae usually have three parts, the left domain that included the hyper variable segment 1 (HVS-1) and a tandem repeated region in the 5’ end, the central conserved region (CCR) and the right domain that included the hyper variable segment 2 (HVS-2) and a tandem repeated region in the 3’ end ^12^. We did not find tandem repeats in the left domain (15,460-15,737 bp) of the *S. populator* mtDNA control region (which may be due to short read sequencing bias), however, like most Felidae, the 3’ end of the right domain (16,261-16,769 bp) of this species presented a tandem repeated region, with a repeat unit of 41 bp repeated five times (3’-CACCCACGTGTACGTATACGCGCACAGCACGCACACACGTA - 5’). Unlike *S. populator*, the left domain (15,468–16,332) of the Control Region of *H. latidens* had a tandem repeated region with an 80 bp repeat unit repeated 7.3 times (3’-ATAAAACATACTATGTATATCGTGCATTAACTGCTAGTCCCCATGAATAATAAGCATGTACCATACATTCATTATAATAC-5’) and the right domain (16,906-17,323 bp) presented an 8 bp repeat unit repeated 14 times (3’-CGTATACG-5’). This characteristic makes the *H. latidens* D-loop very similar to other cats mitogenomes, like the Asian lion ^12^. As expected, for both species the most common start codon was ATG, however in both species the ND2 gene starts with ATT and the *ND5* gene starts with ATA. Most of the felids *ND2* gene starts with the ATC codon, although a few cases like *Acyonyx jubatus* and *Catopuma temminckii* starts with the ATT codon like *S. populator* and *H. latidens* ^10,13^. Interestingly, except for Hyaenidae, represented in our analysis by *Crocuta crocuta* and *Hyaena hyaena*, all outgroups also presented ATT as start codon in the *ND2* gene. For *ND5* gene, however, like *S. populator* and *H. latidens* most of the felids and outgroup species included in the phylogeny had ATA as a start codon, except for *Viverricula indica* and *Genetta servalina* which starts with ATT. Both species had also the same stop codons for all PCGs, being TAA for *COXI, COXII, ATP8, ND4L, ND5* and *ND6*; TA-for *ND1* and *ATP6*; T--for *ND2, COXIII* and *ND4*; and AGA for *CYTB*. These stop codons are common in felids mitogenomes, with few variations in general related to the presence/absence of the 2^nd^ and 3^rd^ nucleotide on codon TAA/TAG ^10,11^. The relative synonymous codon usage analysis (RSCU) reveals that both species had very similar patterns of RSCU. Both species also showed a preference for A and C bases in third positions of codons, with exception of Tyr and Ile amino acids that presented a preference for U instead C, and a notable avoidance of the G nucleotide in the last position of the codons (Figure2). For both species, the most used amino acid was Leucine (15.69% for *S. populator* and 15.56% for *H. latidens*), followed by Isoleucine (8.75% for *S. populator* and 8.91% for *H. latidens*) and Threonine (8.59% for *S. populator* and 8.15% for *H. latidens*). In both, the least used amino acid was Cysteine (0.58% for *S. populator* and 0.63% for *H. latidens*).

**Figure2.**
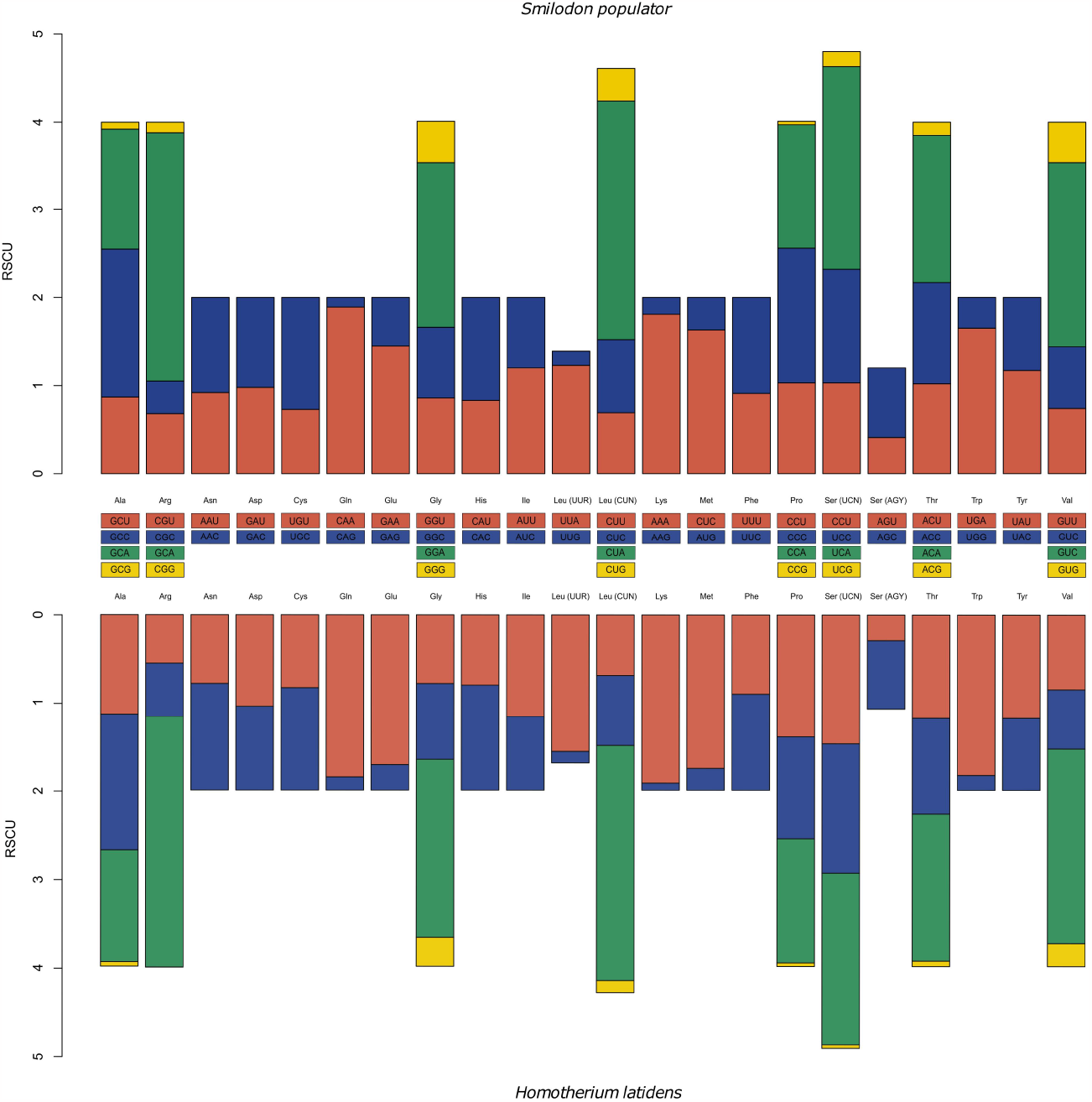
RSCU in *Smilodon populator* and *Homotherium latidens* mitogenomes.

### 2.2 Mitochondrial phylogeny

Both the maximum likelihood and Bayesian trees had the same topology, with in general high values of bootstrap and posteriori probability (Figure3). We recovered the three Felidae subfamilies included in the analysis (Machairodontinae, Pantherinae and Felinae) as monophyletic groups, with Machairodontinae being a sister group of living felids subfamilies, in the same way as previous studies ^1,7^. North American *Homotherium* specimens (SP1714 and YG 439.38) were paraphyletic in relation to the European specimen (SP1007), same phenomenon observed by Paijmans et al. ^1^. It’s unclear if the group of SP1007 and SP1714 specimens are biased due to the assembly method of these mitogenomes (mapping against the YG 439.38 specimen mitogenome). However, the proposal that all Late Pleistocene Holarctic specimens of *Homotherium* are the same species is not new and is based on both morphological and genetic data ^1,5,6^.

**Figure3.**
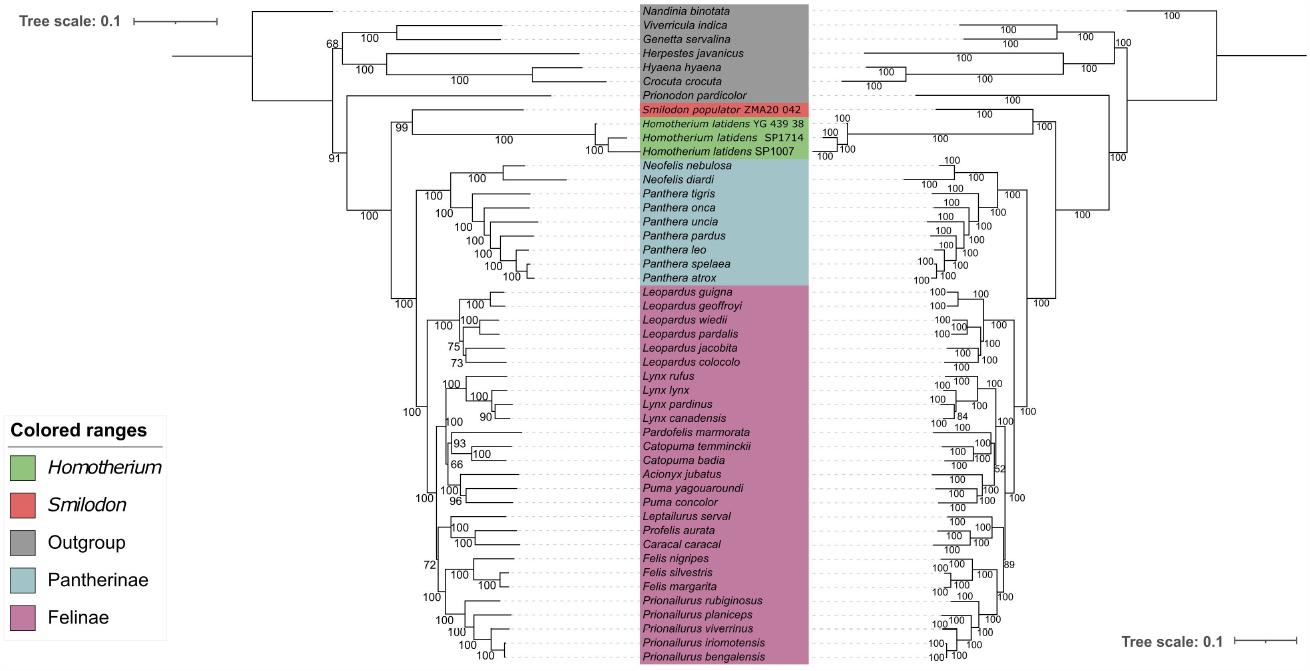
Maximum likelihood (left) and Bayesian (right) trees.

Different phylogeographic and demographic scenarios were proposed to explain the similarity between North American and Eurasian specimens of *Homotherium* from Late Pleistocene. Several species of *Homotherium* from Eurasian Middle Pleistocene were proposed, and some studies have proposed that all of them belongs to *H. latidens* species ^6^. However, Late Pleistocene records in Europe are extremely rare compared with North America, with the most notable case being the North Sea mandible SP1007, a specimen dated to 28 ka while other European *Homotherium* remains are dated until 400-300 ka ^14^. Antón et al. ^6^ pointed out that *H. latidens* were a morphologically variable species, and even the Late Pleistocene specimens from North American, traditionally assigned to *H. serum*, fall inside the *H. latidens* morphological variation. So, the presence of the North Sea mandible SP1007 appears to reaffirm a possible re-colonization of Europe from North America or from some population of Central Eurasia or Beringia ^1,6^. A demographic explanation for the low abundance of Late Pleistocene *Homotherium* fossils is that this taxon presented low population densities, being below the “fossil detection threshold”, but was a highly mobile taxon with a widespread distribution in Eurasia ^1^. However nuclear genome analysis of *H. latidens* has indicated high levels of genetic diversity and heterozygosity suggesting that it was an abundant taxon in Late Pleistocene ^8^.

Unfortunately, due to the absence of other *Smilodon* mitochondrial DNA available, we are unable to bring great advances in the understanding of phylogeny and evolution in this taxon now. However, our complete assembly of *S. populator* mitogenome can be very useful as new mitogenomes, or even single mitochondrial genes, from *Smilodon* or other Machairodontinae genera become available, especially considering the existence of another Late Pleistocene species of *Smilodon*, the *S. fatalis* ^1,2^. Both species traditionally were considered allopatric sister species separates by the Andes, however an almost complete skull of *S. fatalis* found in Uruguay challenged this view, bringing the need to not only review the biogeographical history of the group, but also the taxonomic status of various materials assigned to *S. populator* in the east of the Andes South America ^3^.

### 2.3 Time-calibrated mitochondrial phylogeny

Our chronogram (Figure4) estimated the median time to the most common ancestor (tMCA) of the tree Felidae subfamilies (Felinae, Pantherinae and Machairodontinae) at 21.84ma (95% HPD: 27.578-16.232ma), which was similar to Paijmans et al. ^1^ estimative. For the living Felidae subfamilies (Pantherinae and Felinae), the median time to tMCA was estimated at 16.092ma (95% HPD: 20.051-12.318ma), which was inside the range of previous estimates ^1,15^. For the split between *Smilodon* and *Homotherium* the median time to tMCA was 17.151ma (95% HPD: 23.785-10.858ma) reinforcing an Early Miocene divergence for its groups ^1,16^. The main difference between our Molecular Clock to the Paijmans et al. ^1^ is the age of tMCA between the three *H. latidens* specimens, while Paijmans et al. ^1^recovered a Late Pleistocene (215.970-77.076 ka) divergence to these specimens, we recovered a deep origin for their tMCA (median of 1.68 ma, 95% HPD: 2.317-1.15ma).

**Figure4.**
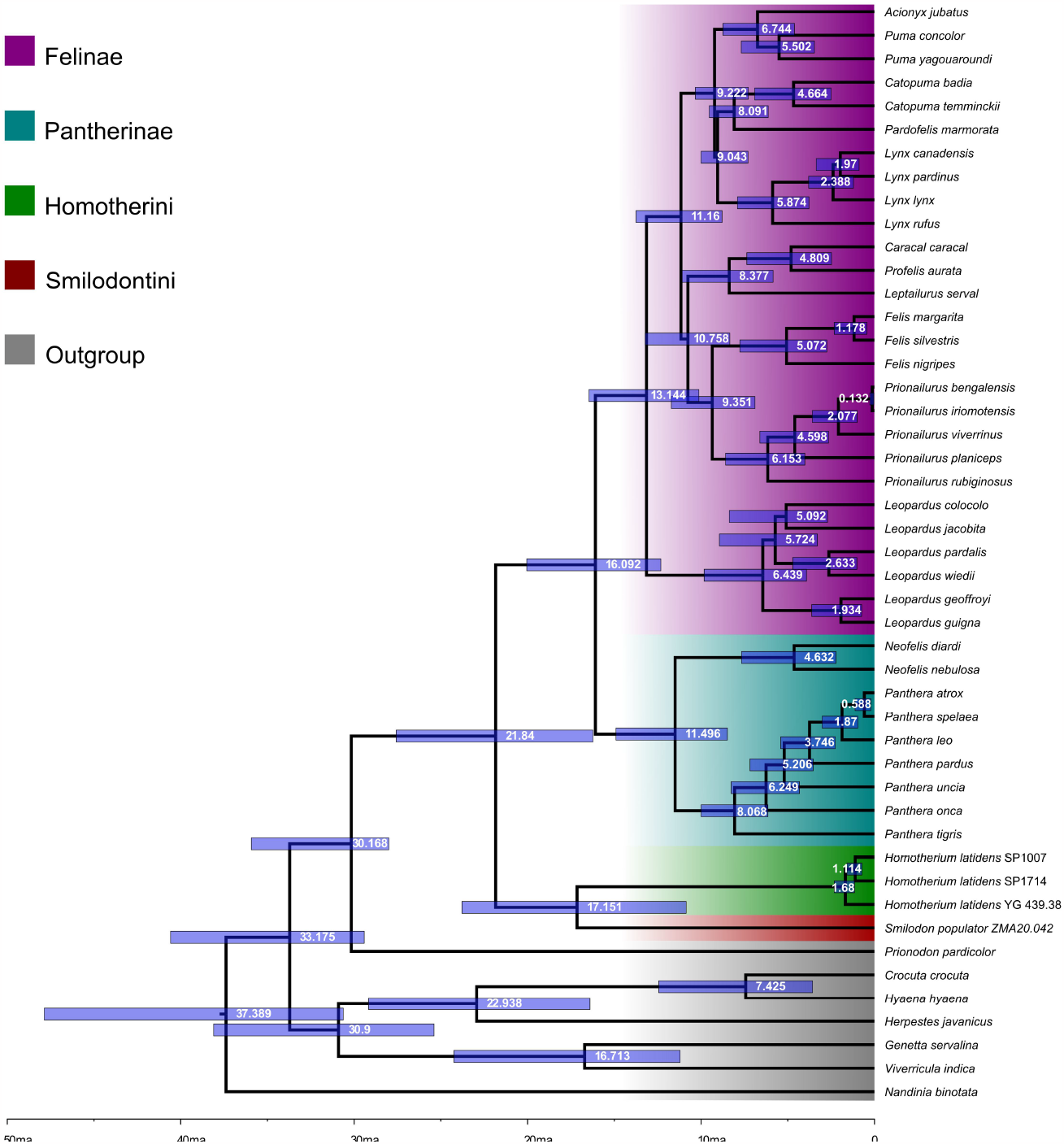
Time-calibrated Bayesian tree.

North American *H. latidens* (aka *H. serum*) are commonly assigned as a Late Pleistocene taxon, although some specimens earlier this age were reported, ^17^*Homotherium* specimens for the Pliocene-Early Pleistocene North America (aka Blancan North American Stage) are generally assigned to older species *Homotherium ischyurus* ^6^. However, it is known that *H. latidens* roamed the Eurasia at this age (Villafranchian age), with one of the oldest species, the “Perrier skull”, dated about 3.3ma ^6^. The two splits (1.68 ma - 95% HPD: 2.317-1.15 ma and 1.114ma – 95% HPD: 1.594-0.722 ma) between the *Homotherium* specimens in our chronogram matches with glacial deposits attributed to pre-Illinoian stage glaciations ^18^. The Early Pleistocene was characterized by symmetric glacial cycles of 41 thousand years separated by shorts interglacial periods ^19^, these glacial periods were characterized by low sea levels, advance of continental ice sheets and by the connection of North America and Eurasia via Beringia. It’s possible that *H. latidens* crossed into Beringia at this time, as has been reported for mammoths ^20,21^.

## 3 Conclusion

Our complete assemblies of *H. latidens* and *S. populator* mitochondrial genomes by a *de novo* method, brings new knowledge of the composition, organization, and peculiarities of the mitogenomes in these extinct taxa. Besides having the same gene order and composition of extant cats and sharing most part of start/stop codons, we observe that the mitochondrial genomes of these species had a slightly lower GC content than modern cats (that in general has more than 39% of GC content) and an unusual start codon for *ND2* gene in cats but that is common in other Feliformia taxa. The two mitogenomes are also very similar to each other, considering gene size and RSCU, with most differences concentrated on the D-loop, with *Homotherium* having a D-loop more similar to extant cats than *Smilodon*. Finally, our analysis reveals that the divergence between the North American and North Sea *H. latidens* could be earliest than expected. We also concluded that, once paleontology and genomics are getting closer every day, the improvement of methods for assembling, annotating and comparison of genomic sequences has great potential in solving or help with phylogenetic and evolutionary issues faced by paleontologists and evolutionary biologists, in addition to help the understanding of genomic dynamics and evolution for extinct and living taxa.

## 4 Material and Methods

### 4.1 Mitochondrial assembly and annotation

We download the raw data of *S. populator* ZMA20.042 (SRR13403298: 17G bases) and *H. latidens* YG 439.38 (SRR12354130: 15.8G bases) from NCBI SRA. To obtain the mitogenome we assemble the raw reads in plasmid-referring contigs with plasmidSPAdes tool v.3.15.4 ^22^, since mitochondrial genomes are extranuclear elements, with high coverage per cell and with bacterial origin. Then we filtered the scaffolds obtained by the expected size of the mitogenomes (15 – 18 kbp) and made a blast search against the NCBI’s nucleotide database to search for contigs that aligned with Felidae mitogenomes.

We annotated the mitogenomes with the MitoAnnotator tool ^9^ at the MitoFish v.3.91 server (http://mitofish.aori.u-tokyo.ac.jp). This tool automatically finds the *tRNA-Phe* gene and adjust the mitogenome coordinates to start with this gene, besides annotating all standard genes typically found on vertebrates mitogenomes. Besides having a database focused on fish mitogenomes, the MitoAnnotator tool works very well with most Vertebrates mitogenomes due to their stable organization and composition. However, to avoid possible biases we also confirm our annotations with MitoZ ^23^ and MITOS2 v.2.1.3 ^24^ tools. We searched for tandem repeated regions on control region for both mitogenomes with the Tandem Repeats Finder v.4.09 ^25^ and calculated the relative synonymous codon usage (RSCU) in the Mega 11 software ^26^.

### 4.2 Mitochondrial mapping of *H. latidens* SP1007 and SP1714

Since we failed to assembly the mitogenomes of the libraries of the *H. latidens* specimens SP1007 and SP1714 by *de novo* method, we decided to recover the mitochondrial sequences of these specimens by mapping the libraries against our *H. latidens* YG 439.38 mitochondrial genome assembly. First, we import all libraries for these two specimens (SRR1970776, SRR1970777, SRR1970778 and SRR1970779 for SP1007; SRR1970780, SRR1970781, SRR1970782 and SRR1970783 for SP1714) into the Galaxy Europe platform ^27^, then we concatenate all libraries per specimens and used the fastp tool v.0.23.2 ^28^ to trimming the reads with q>=30 and discarding reads with length shorter than 30 nt. We used the Bowtie2 ^29^ tool v.2.5.0 to map the concatenated libraries against complete mitochondrial genome and all separately mtDNA genes of *H. latidens* YG 439.48. After, we used the ivar consensus tool v.1.4.2 ^30^ to obtain the consensus sequences in bowtie alignments. Then, we downloaded the fasta sequences and aligned the mapped genes against the mapped mitogenome contig in Mega 11 software ^26^. Finally, we obtained the consensus sequence of the mitogenomes with bioseq package ^31^.

### 4.3 Mitogenome comparisons with BRIG

To compare our assemblies against the partial mitochondrial genomes available at the NCBI’s database, we conducted a comparative BLAST analysis with the BLAST Ring Image Generator (BRIG) tool ^32^ with published assemblies of the specimens *S. populator* ZMA20.042 (MF871700.1) and *H. latidens* YG 439.38 (MF871702.1) using our assemblies of the same specimens as reference. For the two *H. latidens* assemblies by mapping method, we used our *H. latidens* YG 439.38 as reference and aligned our mitogenomes and the published ones of SP1007 (MF871701.1) and SP1714 (MF871703.1) specimens.

### 4.4 Mitochondrial DNA phylogeny

For the phylogeny, we extracted the 13 PCGs and the 2 rRNAs of our four assemblies and downloaded from NCBI’s nt database the same genes from other 36 felids mitogenomes plus seven outgroups of the families Hyaenidae, Herpestidae, Prionodontidae, Viverridae and Nandiniidae. We aligned each gene with Mafft tool v.7.508 ^33^ and removed from alignment’s sites with gaps in at least 10% of taxa. We used the Concatenator tool v.0.2.1 ^34^ to concatenate the alignments, generate partitions by each gene and codon (for PCGs only) and inputs for IQ-TREE and MrBayes software’s partitioned for each gene. We perform a maximum likelihood tree in IQ-TREE v.2.1.2 software ^35^ with 10000 ultrafast bootstrap replications and a Bayesian inference tree in MrBayes v.3.2.7 ^36^ with 10 million generations, sampling every 100 generations, four chains and two runs, with burnin set in 25%. For the maximum likelihood, the evolutionary models were calculated by the ModelFinder ^37^ of IQ-TREE and for Bayesian inference, we used the PartitionFinder v.2.1.1 ^38^ to obtain the best evolutionary model available on MrBayes. We rooted the trees in *Nandinia binotata*.

### 4.5 Molecular clock

We estimated divergence times for the group phylogeny based on a time-calibrated Bayesian analysis using BEAST v.2.6.6 software ^39^. We run the analysis with a lognormal clock model, using as tree parameter the Birth-Death model. For fossil calibration we followed the calibrations priors of Paijmans ^1^ and used the oldest *Genetta* fossil to limit the split between *Genetta servalina* and *Viverricula indica* with an uniform prior of 50-11.2ma; the oldest herpestid and hyaenid fossils to limit the split between Herpestidae and Hyaenidae with an uniform prior of 50-16.4ma; the stem felids and *Prionodon* fossils to limit the split between *Prionodon pardicolor* and Felidae with an uniform prior of 50-28ma; the oldest *Lynx* fossil to limit the common ancestor of *Lynx, Pardofelis* and *Catopuma* genera with an uniform prior of 10-5.3ma; the oldest *Acinonyx* fossil to limit the common ancestor of *Puma yagouroundi, Puma concolor* and *Acinonyx jubatus* with an uniform prior of 10-3.8ma; the oldest *Caracal* and *Leptailurus* fossils to limit the common ancestor of *Leptailurus serval, Profelis aurata* and *Caracal caracal* with an uniform prior of 16-3.8ma; the oldest *Panthera* fossil to limit the split between *Neofelis* spp. and *Panthera* spp. with an uniform prior of 16-3.8ma; and the oldest *Panthera tigris* fossil to limit the clade containing all *Panthera* spp. with an uniform prior of 10-1.5ma. We perform two independent MCMC chains, with 50 million generations and tree sampling every 5,000 generations. As nucleotide substitution model we chose for each partition the best evolutionary according to the PartitionFinder v.2.1.1 ^38^ analysis, however, since some partitions failed to achieve convergence with the GTR+G+I model, we change the substitution model for these partitions to HKY+G+I. We used Tracer v.1.7.1 ^40^ to check the convergence and posterior samples of the MCMC chain parameters (ESS>200). The tree files were combined using LogCombiner v.2.6.6 ^39^ with burn-in in 10% and we used the TreeAnnotator v.2.6.6 ^39^ to reconstruct the chronogram.

## 5 Declarations

The authors would like to thank the financial support provided by Capes, Fapemig and CNPq through the scholarships granted to IHRO, IBS, RRR and RASS. The complete mitochondrial genome sequences have been submitted to genbank and are under analysis, accession numbers will be provided as soon as possible.

